# Evolution of the cellular landscape in mammalian striatum

**DOI:** 10.1101/2025.04.20.649707

**Authors:** Gozde Buyukkahraman, Emre Caglayan, Stephen G. Hoerpel, Yaqiang Zhang, Ine A. van Tussenbroek, Carlos G. Orozco, Emily Oh, William D. Hopkins, Chet C. Sherwood, Todd F. Roberts, Sonja C. Vernes, Genevieve Konopka

## Abstract

The dorsal striatum is important for highly specialized functions including movement, learning, and habit formation. However, it is not known if species-specialized behaviors are associated with cellular specializations in the striatum. Here, we compared single-nucleus RNA sequencing (snRNA-seq) data from human, chimpanzee, rhesus macaque, common marmoset, and pale spear-nosed bat caudate (CN) and putamen (Pu) separately as well as mouse caudoputamen (C-Pu), which represents divergence among species spanning approximately 94 million years of evolution. We observed a lower neuron-to-glia ratio in primate striata compared to non-primates, reflecting the allometric scaling of neuron density and relative glia density invariance in larger brains. Among neurons, eccentric spiny projection neurons (eSPNs) - an SPN of unknown function - showed significantly lower proportions in non-primate striata for both CN and Pu. Focusing on the heterogeneity within interneurons, we identified two bat striatal interneuron cell types that are nearly absent in other species: which express *LMO3*, and co-express *FOXP2* and *TSHZ2*. Other striatal interneurons also showed significantly differential abundance between primates and non-primates. In summary, we provide a comprehensive snRNA-seq dataset of dorsal striatum, identify novel interneuron innovations in bat, and uncover fundamental cellular composition differences between primate and non-primate striata.

## Introduction

Comparing the cellular composition of the striatum among humans, primates and other mammals is compelling because it makes vital functional contributions to behaviors such as decision making (Cox and Witten 2019), learning (Cataldi, Stanley et al. 2022), and (vocal) motor control (Volkmann, Hefter et al. 1992, Tressler, Schwartz et al. 2011, Etchell, Johnson et al. 2014, Watkins 2016, Weineck, Garcia-Rosales et al. 2020), as well as habit formation (Yin, Knowlton et al. 2004, Graybiel and Grafton 2015, Phan, Ray et al. 2024) and goal-directed behavior (Li, Yan et al. 2008, Schel, Ridderinkhof et al. 2014, Yoshida and Hikosaka 2024). The striatum receives most of its input from the cerebral cortex and provides feedback via pallido-thalamic circuits, making the study of striatal cell types relevant to building cell type-based circuit molecular models (Alexander, DeLong et al. 1986). The striatum is also a major interface between cortical regions and subcortical structures in the control of behavior (Macpherson, Morita et al. 2014). Not surprisingly, the disruption and dysfunction of striatal circuits lead to many devastating diseases such as Parkinson’s disease (Zhai, Tanimura et al. 2018), Huntington’s disease (Vonsattel, Myers et al. 1985, Hodges, Strand et al. 2006, Malaiya, Cortes-Gutierrez et al. 2021, Matsushima, Pineda et al. 2023), Tourette’s syndrome (Albin, Koeppe et al. 2003, Hienert, Gryglewski et al. 2018) and obsessive-compulsory disorder (OCD) (Graybiel and Rauch 2000, Ahmari, Spellman et al. 2013, Gillan, Morein-Zamir et al. 2014, Gillan and Robbins 2014), thus, making it a key area of study for understanding the pathophysiology of these diseases. Furthermore, an evolutionary perspective is vital for understanding the susceptibility of the human brain to neurodegenerative, neurodevelopmental and psychiatric disorders (Diederich, Uchihara et al. 2020, Pattabiraman, Muchnik et al. 2020). While much progress has been made in comparing the human brain to a limited number of model organisms, the majority of such studies have examined only a few different regions of comparison, primarily within the cerebral cortex (Khrameeva, Kurochkin et al. 2020, Ma, Skarica et al. 2022, Caglayan, Ayhan et al. 2023). A recent study comparing cortical and striatal interneurons between primates and rodents found a transcriptionally distinct interneuron subtype only in the striatum, with no such observations in the cortex (Krienen, Goldman et al. 2020), highlighting the necessity of further cellular resolution comparative investigations of the striatum.

The dorsal striatum is comprised of the caudate nucleus (CN) and putamen (Pu), two morphologically distinct brain regions separated by the internal capsule in carnivores and primates (Mink 2013, Ni, Huang et al. 2018). The CN and Pu can be functionally distinguished in human brain imaging studies during certain tasks such as those involving language or speech (Tettamanti, Moro et al. 2005, Booth, Wood et al. 2007). Within CN and Pu, GABAergic spiny projection neurons (SPNs) comprise the majority of neurons (Gerfen, Engber et al. 1990, Day, Belal et al. 2024). The SPNs themselves form two major subtypes. The dopamine receptor 1 (DRD1) expressing SPNs (dSPNs) form the direct pathway and project to the globus pallidus internus (GPi) and the substantia nigra pars reticulata (SNr). The dopamine receptor 2 (DRD2) expressing SPNs (iSPNs) form the indirect pathway and project to the globus pallidus externus (GPe), which subsequently projects to the subthalamic nucleus (STN) (Alexander and Crutcher 1990, Gerfen, Engber et al. 1990). In addition to the two subtypes, a third class of SPNs, eccentric SPNs (eSPNs), has been identified and defined by transcriptomic signatures rather than classical DRD1/2 expression (Saunders, Macosko et al. 2018, Anderson, Kulkarni et al. 2020). Previous studies have identified and described the function of SPNs that express both D1 and D2 (D1/D2 hybrid SPNs) in rodents (Aizman, Brismar et al. 2000, Shuen, Chen et al. 2008, Matamales, Bertran-Gonzalez et al. 2009, Thibault, Loustalot et al. 2013, Bonnavion, Varin et al. 2024) and primates (Aubert, Ghorayeb et al. 2000, He, Kleyman et al. 2021, Phan, Ray et al. 2024); however, it remains to be determined whether these D1/D2 hybrid cells can be characterized as eSPNs.

Interneurons, identified by the neuropeptides that they express, make up the rest of the striatal neurons and regulate the cell signaling, excitability, and spiking timing of SPNs (Tepper, Tecuapetla et al. 2010, Burke, Rotstein et al. 2017). A few studies have begun to compare the cellular composition and cell type expression patterns of the human striatal interneurons with that of other species. For example, Krienen et al identified a primate-specific striatal interneuron cell type, defined by expression of *TAC3,* which had no homologous molecular counterpart in non-primate striata (Krienen, Goldman et al. 2020). Whereas a recent study that expanded the Boreoeutherian mammal species sampled to include ferrets and pigs, showed that the initial classes forming *TAC3* interneurons are conserved across 90 million years of evolution (Corrigan, DeBerardine et al. 2024). Using single nuclear RNA sequencing (snRNA-seq) and spatial transcriptomics, a detailed classification of human striatal interneurons revealed that *PTHLH* and *TAC3* interneurons represent the largest classes of interneuron cell types with the CN showing a significant abundance of the *PTHLH* subtype compared to the Pu (Garma, Harder et al. 2024). These studies provide the initial classification of striatal interneurons and reveal their remarkable heterogeneity that has been subjected to evolutionary change. However, a comprehensive comparison of human striatum to a diverse group of species has yet to be performed.

In this study, to assess the extent of conservation in terms of cell types, and gene expression profiles in the dorsal striatum (called “striatum” hereafter) across functionally distinct regions, we generated snRNA-seq datasets of CN and Pu. To make comparisons among primate species, we included data from adult humans, chimpanzees, rhesus macaques, and common marmosets. We found the proportion of neurons was significantly higher in the CN of macaque and marmoset compared to human and chimpanzee, but these differences were not observed in the Pu. To extend these findings to non-primate mammals we included adult mouse striatum dataset (Krienen, Goldman et al. 2020) and to include a non-primate mammal with distinct CN and Pu structures, we generated CN and Pu snRNA-seq datasets from adult pale spear-nosed bats. We found a strikingly lower neuron-to-glia ratio in primate striata compared to non-primates. By including CN and Pu datasets, we were able to identify cellular features that are conserved across mammalian CN or Pu, as well as features that are unique to pale spear-nosed bats. Together, these findings provide comprehensive cellular and transcriptomic comparisons across mammalian CN and Pu while simultaneously providing the first bat and chimpanzee striatum snRNA-seq datasets.

## Results

### Identification of striatal cell populations of each species

To understand cellular evolution of the dorsal striatum across diverse mammal species, we generated snRNA-seq data from caudate nucleus (CN) and putamen (Pu) postmortem brain tissues from adult human (CN n=6 and Pu n=7), chimpanzee (CN n=4 and Pu n=6), rhesus macaque (CN n=4 and Pu n=7), and pale spear-nosed bat (CN n=4 and Pu n=4) (**Fig. 1A, Fig. S1A**). In addition to these newly generated datasets, to extend our species comparisons, we included published snRNA-seq datasets from adult rhesus macaque (He, Kleyman et al. 2021), common marmoset (Lin, Kelly et al. 2022, Krienen, Levandowski et al. 2023), and mouse (Krienen, Goldman et al. 2020) (**Fig. 1B)**.

**Fig. 1:**
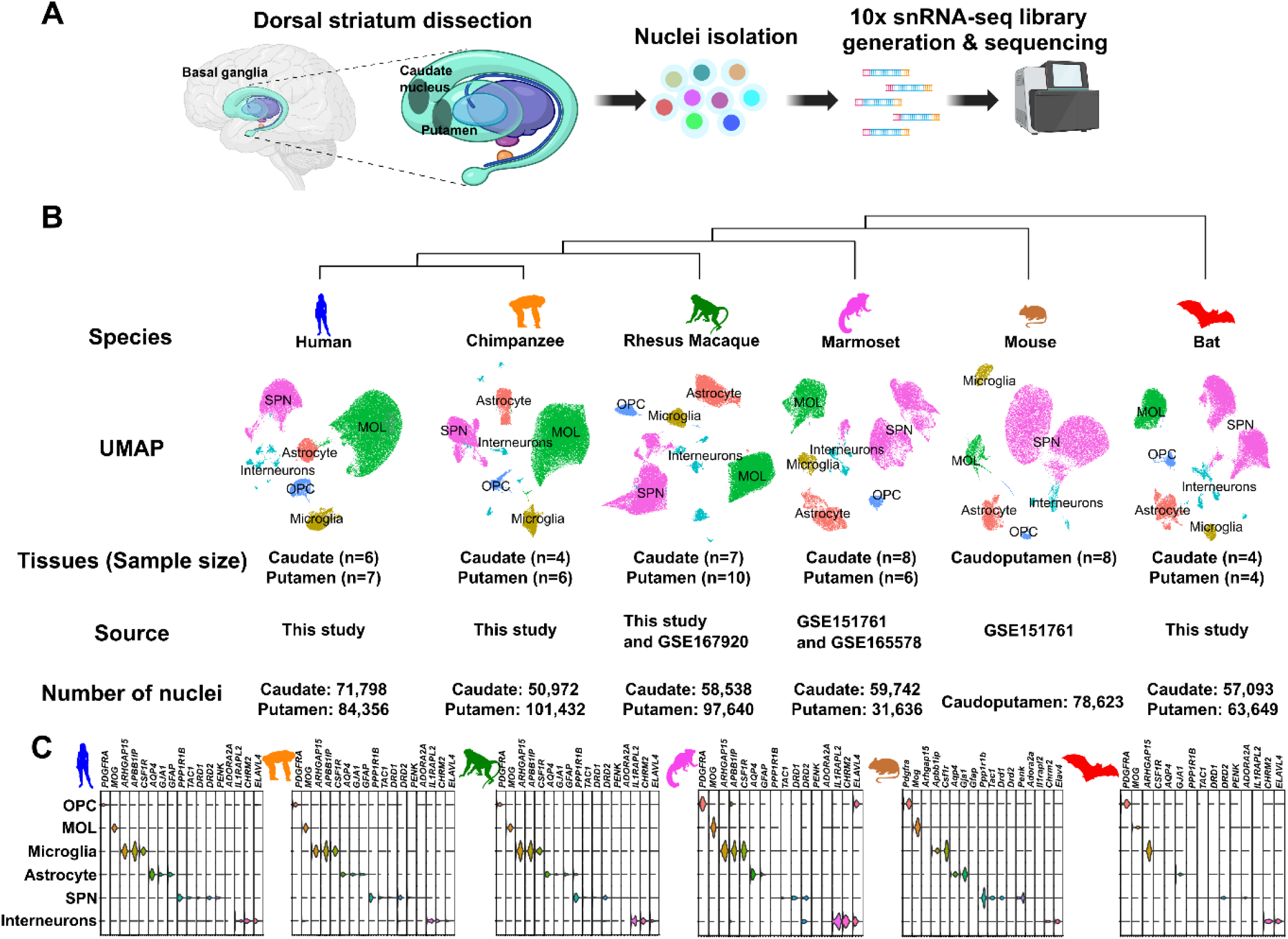
Overview of the snRNA-seq datasets. **A.** Workflow depicting the dissection of the caudate nucleus (CN) and putamen (Pu) and the subsequent preparation of snRNA-seq libraries. **B.** Summary of species, major cell type annotations, sample sizes, sources, and the number of nuclei post-filtering for the snRNA-seq datasets analyzed in this study. **C.** Violin plots illustrating the expression of marker genes for major cell type annotations across different species. Animal illustrations were obtained from https://www.phylopic.org/. Panel A illustrations were created with BioRender.com.

We integrated all datasets from raw reads and implemented consistent quality control to both the new and published datasets (see **Methods**). After stringent quality control, we identified 156,154 cells in human, 152,404 cells in chimpanzee, 156,178 cells in rhesus macaque, 91,378 cells in marmoset, 78,623 cells in mouse, and 120,742 cells in bat (**Fig. 1**). The mean number of unique molecular identifiers (UMIs) of the samples we sequenced was at or above 1000, comparable to the published datasets (**Fig. S1B**). To evaluate the quality of the nuclei, we further employed intronic read ratio, a reliable quality control metric for snRNA-seq data (Caglayan, Liu et al. 2022), and found that intronic read ratios were consistently high across all samples (**Fig. S1C**).

We identified major cell types in all species using the conserved cell type markers: *AQP4*, *GJA1* and *GFAP* for astrocytes, *CSF1R*, *ARHGAP15*, and *APBB1IP* for microglia, *MOG* for mature oligodendrocytes (MOLs), *PDGFRA* for oligodendrocyte progenitor cells (OPCs), *PPP1R1B*, *DRD1, TAC1, PENK, and DRD2* for SPNs, and *IL1RAPL2, CHRM2,* and *ELAVL4* for interneurons (see **Methods, Fig. 1C and Fig. S1D**). While we observed similar contributions to cell types across most samples within species, there were striking differences of cell type proportions across species as well as within species between CN and Pu (**Fig. S1E-F**), which motivated us to examine them further.

### Proportional differences in neuron and glia across species

To determine cell type compositional differences, we first calculated the proportion of neurons (number of neurons divided by total number of cells, neurons and glia) in each combined (CN + Pu) dataset and compared these values across species. Human samples had a mean neuron proportion of 26.2%. Interestingly, this ratio increased with increasing evolutionary distance from humans and decreasing brain size (**Table S2** and **Fig. 2A**). While neuronal proportions were not significantly different between human and chimpanzee, both macaque and marmoset neuronal proportions were significantly greater than human (two-tailed t-test p = 0.0073 and p = 2.3 × 10^-4^, respectively. **Table S2** and **Fig. 2A**). As an outgroup to the primates, we calculated neuronal proportions in mouse and found that they were significantly higher than humans (two-tailed t-test p=0.0018). Since mouse does not have distinct CN and Pu structures but instead has a caudoputamen (C-Pu) structure, we included another non-primate, bat, where we can distinguish these structures. We calculated the neuronal proportions in bat and found that similar to the other species, bat has higher levels of neuronal proportions than human (two-tailed t-test p = 9 × 10^-10^), although bat neuronal proportions displayed high variation (59.5% ± 20.3%) (**Table S2** and **Fig. 2A**). These results indicate a significantly lower neuron-to-glia ratio in primates, which we interpret to correlate with larger brain size compared to the non-primates in the dataset.

**Fig. 2:**
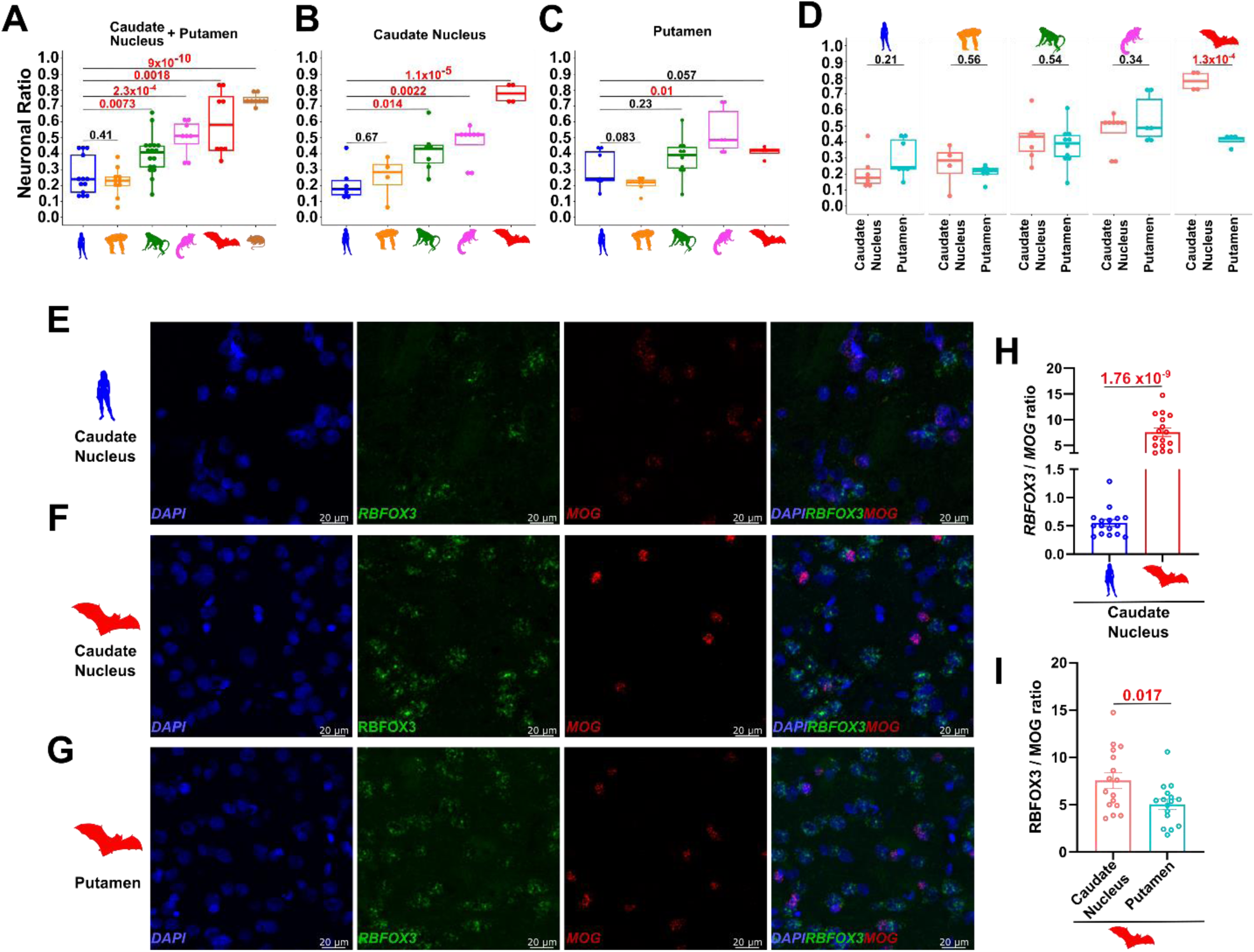
Neuronal ratios across species. Ratios were calculated by the number of neurons (SPNs and interneurons) divided by the number of total cells (neurons and glia) within each species. We compared each species with human in the following tissue: **A**. Caudate Nucleus (CN) and Putamen (Pu) (Dorsal Striatum), **B**. CN, and **C**. Pu. **D**. Neuronal proportions were compared between CN and Pu within each species. smFISH validation of **E**. human CN, **F**. bat CN, and **G.** bat Pu. The scale bars represent 20 µm. **H.** The ratios of the number of cells co-stained with *RBFOX3* and DAPI and the number of cells co-stained with *MOG* and DAPI in human and bat CN (n=16 each) are shown. **I.** The ratios of the number of cells co-stained with *RBFOX3* and DAPI and the number of cells co-stained with *MOG* and DAPI in bat CN and Pu (n=16 each) are shown. Two tailed t-test was used to assess statistical significance between the groups. For comparisons where statistical significance was observed, p-values are highlighted in red. Error bars indicate mean ± standard error of the mean.

A recent study showed that there are significant differences in striatal interneuron cell type proportions between CN and Pu in human striatum (Garma, Harder et al. 2024). However, a comprehensive analysis of the cellular composition, encompassing all cell types in these striatal regions, has not been conducted in detail across multiple species. To investigate potential regional contributions to interspecies variability in neuronal proportions (**Fig. 2A**), we compared the proportions of neurons in the CN and putamen Pu across species separately. We excluded mice for these comparisons as their C-Pu structure does not morphologically distinguish between CN and Pu. We found similar evolutionarily divergent neuronal proportions in CN but not in Pu, indicating that combined striatal results are primarily driven by the CN samples (**Fig. 2B-C**). Bat samples were particularly notable as neuronal proportions were ∼40% in Pu compared to ∼80% in CN (**Fig. 2B-C**). Comparing neuronal proportions between the CN and Pu within each species, we found that only bat displayed a drastic and significant change with a greater neuronal proportion in CN (two-tailed t-test p = 1.3 × 10^-4^, **Fig. 2D**). These results indicate that CN and Pu have similar neuronal cell type proportions within most species, but there may be exceptions in some mammals (e.g. bats in this study) with unexplored consequences for neural circuitry and behavior.

We then asked if lower neuronal proportions in human CN were driven by a relative increase in a subtype of glial cells. To answer this question, we calculated the ratio of neuronal to glial cell number in the CN of each species and in the C-Pu of mouse. We found an overall increase in neuron to glial cell type ratios in other species compared to humans for most glial cell types **(Fig. S2A-D)** with the exception of astrocytes within primates **(Fig. S2C)**. For example, the neuron to MOL ratio was significantly higher in marmoset, macaque and bat CN compared to human CN (**Fig. S2A**). Using intact tissue sections, we independently verified the human and bat comparison in CN using smFISH for *RBFOX3* (neuronal marker) and *MOG* (oligodendrocyte lineage marker) (two-tailed t-test p = 1.1 × 10^-11^) (**Fig. 2E-F, H**).

To further assess which glial cell type proportion have altered neuronal proportions in Pu compared to CN, especially in bats **(Fig. 2D)**, we similarly compared the proportions of neurons to each glial cell type in CN and Pu. We observed that Pu had significantly higher neuron to MOL and neuron to OPC ratios compared to CN only in bats (**Fig. S2E-F**). These differences were further validated through smFISH, confirming significant variation in the neuron-to-MOL ratio between bat CN and Pu (**Fig. 2F-I**). We also observed a trend of higher neuron to astrocyte and neuron to microglia ratios in bat CN compared to bat Pu, although the differences were not significant (**Fig. S2G-H**). The difference was especially low in astrocytes with only a ∼1.5-fold increase of neuron / astrocyte in CN compared to at least 2-fold increase in all other glial cell types **(Fig. S2E-H)**. Given that neuron / astrocyte is also more similar than neuron / other glial cell types across species **(Fig S2A-D)**, this might indicate that astrocyte numbers are more likely to scale with neuron numbers across evolution, a plausible outcome given their critical role in neuronal survival both *in vivo* and *in vitro* (Drukarch, Schepens et al. 1998, Verkhratsky, Butt et al. 2023). Together, these results show that most neuronal proportional differences across species and CN-Pu are not driven by only one glial cell type although the number of astrocyte cells scale more consistently with number of neuronal cells. These findings are congruent with previous observations that neuron density tends to decrease and glial cell density remains invariant with brain size variation in the ventral striatum of primates (Hirter, Miller et al. 2021) and across other brain regions in mammals (Herculano-Houzel 2014).

### Non-primates have lower eSPN to SPN proportions

SPNs comprise the majority of neurons in the striatum, so we identified and annotated the subtypes of SPNs based on the canonical markers *DRD1* and *TAC1* for dSPNs, *DRD2* and *PENK* for iSPNs, and co-expression of dSPN and iSPN markers as well as selective expression of *CASZ1* for eSPNs. Besides containing subtypes of SPNs, the dorsal striatum is patterned with functionally and histochemically different structures called the striosome and matrix, and both structures contain all subtypes of SPNs (Crittenden and Graybiel 2011, Fujiyama, Sohn et al. 2011). While the functional differences of striosome and matrix compartments are yet to be understood, these compartments are differentially affected in disorders associated with basal ganglia dysfunction (Crittenden and Graybiel 2011). We therefore also distinguished which cells represented striosome or matrix by using canonical markers *SPON1*, *BAXH2*, *SEMA5B*, *KREMEN1*, and *OPRM1* for striosomes, and *CALB1*, *CRYM*, *SGK1*, and *SV2B* for matrix (see **Methods, Fig 3A, Fig S3A-B**). We found that the distribution of each sample for these cell types was comparable (**Fig. S3C-D**).

**Fig. 3:**
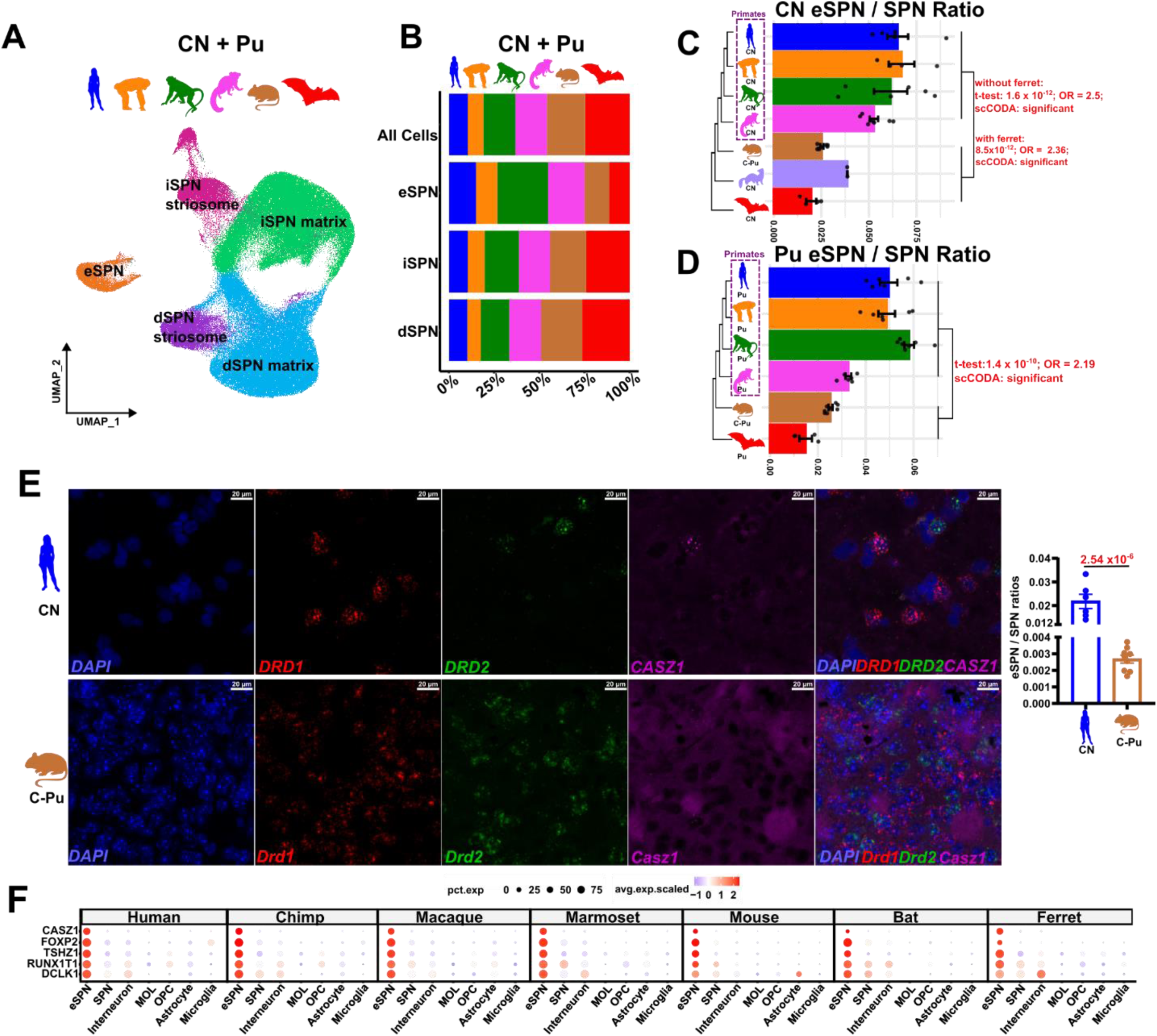
Spiny projection neurons (SPNs) across species. **A**. UMAP of SPNs integrated across human, chimpanzee, rhesus macaque, marmoset, mouse, and bat. **B**. Bar plot depicting the proportional compositions of the species across SPN subtypes. **C**. Bar plot showing the means of the ratios of the number of eSPNs to the total number of SPNs across the mouse caudoputamen, and the ferret, bat, human, chimpanzee, rhesus macaque, marmoset caudate nucleus (CN). **D**. Bar plot showing the means of the ratios of the number of eSPNs to the total number of SPNs across the mouse caudoputamen, and the bat, human, chimpanzee, rhesus macaque, marmoset putamen (Pu). **E.** smFISH validation of human CN and mouse C-Pu. The scale bars represent 20 µm. The ratio was calculated as follows: The number of cells co-stained with *CASZ1, DRD1, and DRD2* and DAPI were divided by the sum of the number of cells co-stained with *DRD1* & DAPI and *DRD2* & DAPI in human CN (n = 6). Similarly for mouse C-Pu, the number of cells co-stained with *Casz1, Drd1, and Drd2* and DAPI were divided by the sum of the number of cells co-stained with *Drd1* & DAPI and *Drd2* & DAPI (n= 9). **F.** Dot plot of eSPN markers across all cell types in human, chimpanzee, rhesus macaque, marmoset, mouse, bat, and ferret. Two tailed t-test and scCODA were used to assess statistical significance between the groups. For comparisons where statistical significance was observed, p-values are highlighted in red. OR: odds ratio. Error bars indicate mean ± standard error of the mean.

The overall composition of SPNs across species suggested that there may be differences in eSPN proportions, especially in bat and mouse (**Fig. 3B**). To test this, we calculated the eSPN to SPN ratios and compared non-primates to primates. We found that the relative proportion of eSPNs among all SPNs was significantly less in non-primates (**Fig. 3C-D**). This difference was significant in both the CN and the Pu (**Fig. 3C-D)**. Additionally, we tested the statistical significance of the comparisons using a more robust proportional analysis of single cell-data (scCODA) to correctly account for the assumptions imposed by the nature of the proportional data (Buttner, Ostner et al. 2021). Using the iSPN cell type as the reference, we compared non-primate eSPN proportions to primate eSPN proportions. We found significantly less eSPN to SPN proportions in non-primates compared to primates, thus confirming the t-test statistics results (**Fig. 3C-D**).

Since mouse has a C-Pu structure and bat has morphologically distinct CN and Pu, we asked if having reduced eSPN to SPN proportions is specific to mouse and bat. To test this, we included another species as an outgroup to primates where CN and Pu can be separated, a ferret CN dataset (Krienen, Goldman et al. 2020) (**Fig. S3E-F)**. We found that high eSPN to SPN proportions were indeed specific to the primates, with non-primates having significantly fewer eSPNs relative to SPNs (p = 8.5 × 10^-12^, OR = 2.36, **Fig. 3C**). We verified these findings using smFISH where human CN showed significantly higher eSPN to SPN ratios compared to mouse C-Pu (two-tailed t-test p = 2.54 × 10^-6^, **Fig. 3E**).

We also examined how similar the eSPNs we identified are to the D1/D2 hybrid cells previously identified in the macaque striatum (He, Kleyman et al. 2021). To achieve this, we performed a correlation analysis of the transcriptional profiles of these cells and found, as previously stated, that they were highly similar (correlation coefficient, r = 0.87), sharing most of the expressed genes and their expression levels (**Fig. S3G)**. Additionally, we found that eSPNs mostly expressed striosome markers (**Fig. S3B**), as previously reported (He, Kleyman et al. 2021). Since eSPNs are relatively understudied, we wanted to identify what other genetic markers might distinguish eSPNs for future functional studies. We thus performed a Wilcoxon rank sum test to find the genes which are differentially expressed by eSPNs compared to other cell types within each species. We then compared these marker genes across the eSPNs of all species. As expected, human eSPN marker genes showed the greatest overlap with chimpanzee and macaque (**Fig. S3H**). The second largest overlap of genes was observed within primates (**Fig. S3H)**. We found *CASZ1*, *FOXP2*, *TSHZ1*, *RUNX1T1*, and *DCLK1* genes mark eSPNs consistently in all species (**Fig 3F**).

### Identification of interneurons found primarily in the bat

Unlike SPNs that project outside the striatum, striatal interneurons largely mediate the local connectivity by regulating the activity of SPNs (Burke, Rotstein et al. 2017). Strikingly, striatal interneurons are very heterogeneous despite making up only 5-10% of the striatal neurons (Gerfen and Surmeier 2011). To uncover striatal interneuron heterogeneity across species, we subsetted and integrated interneurons across species and annotated each cluster based on the expression of the following genes: *CHAT*, *SST*, *NPY*, *PVALB*, *TH*, *TAC3*, *CCK*, *VIP*, *PDGFD*, and *PTHLH* (**Fig. S4A-C**) that resulted in 12 interneuron cell types (**Fig. 4A**). We then hierarchically clustered the interneuron subtypes based on transcriptional profiles to facilitate understanding of potentially similar cell types (**Fig. 4B**). The most similar cell types identified include *CCK VIP-* and *CCK VIP+* interneurons, *PDGFD PTHLH PVALB-* and *PDGFD PTHLH PVALB+* interneurons, as well as *FOXP2 EYA2* and *FOXP2 TSHZ2* interneurons.

**Fig. 4:**
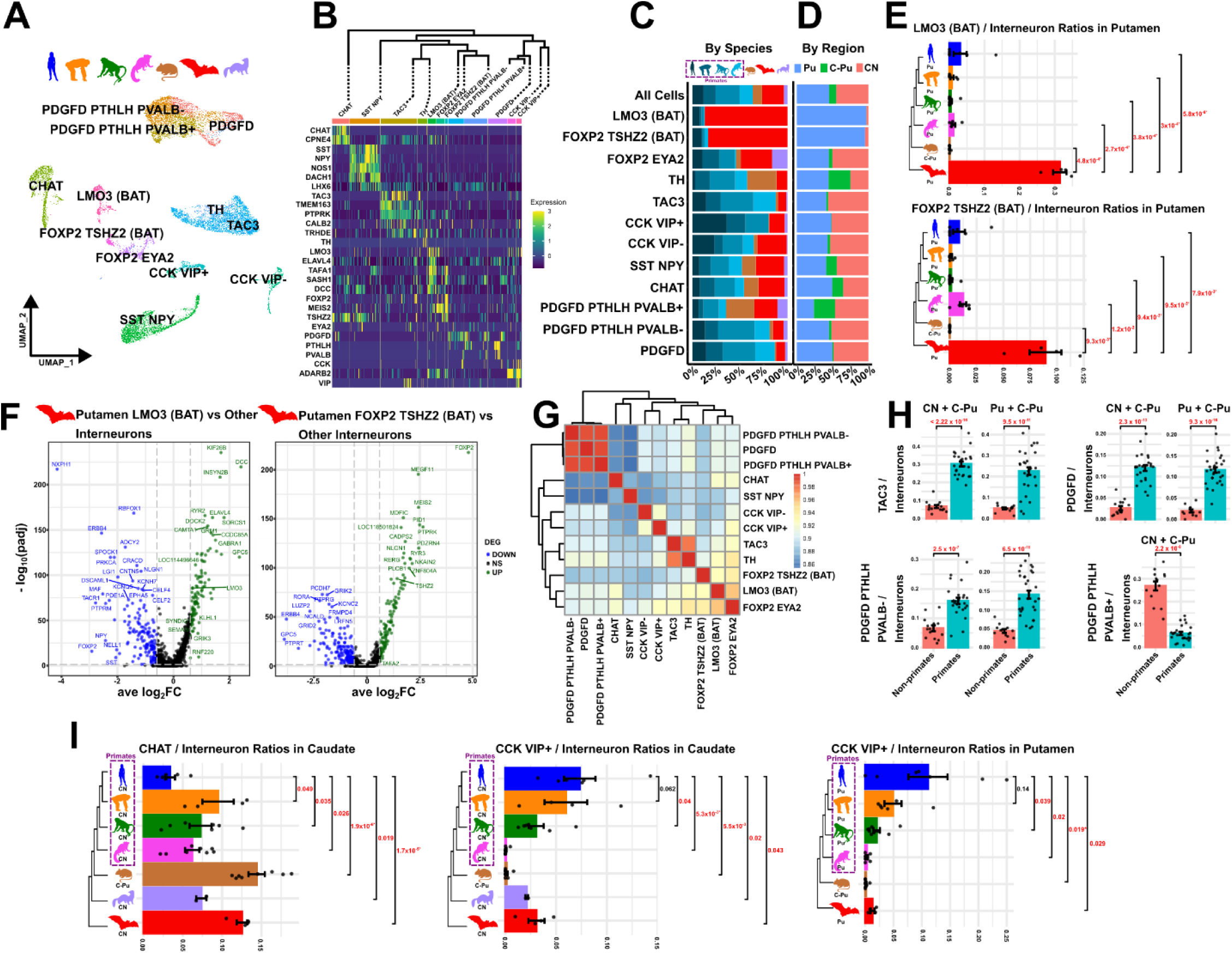
Striatal interneurons across species. **A**. UMAP of interneurons integrated across human, chimpanzee, rhesus macaque, marmoset, mouse, bat, and ferret. **B**. Heatmap showing the gene expression of interneuron cell type markers across the interneuron cell types of all the species. Dendogram shows the hierarchical clustering of the interneuron cell types of all the species. **C**. Bar plot depicting the proportional compositions of the species across interneuron subtypes. **D**. Bar plot depicting the proportional compositions of the regions across interneuron subtypes. Caudate Nucleus (CN), putamen (Pu), and caudoputamen (C-Pu). **E.** Bar plots showing the ratio of the number of *LMO3 (BAT)* and *FOXP2 TSHZ2 (BAT)* interneurons to the number of all interneurons across mouse C-Pu and bat, human, chimpanzee, rhesus macaque, and marmoset Pu. **F.** Volcano plots showing the upregulated and downregulated genes of *LMO3 (BAT)* and *FOXP2 TSHZ2 (BAT)* interneurons compared to interneurons within bat Pu. Green dots represent the significantly upregulated genes (padj < 0.05 and log2FC > 0.6) and blue dots represent the significantly downregulated genes (padj < 0.05 and log2FC < −0.6) **G.** Heatmap showing the correlation of normalized gene expressions across interneurons in bat Pu. Dendrogram depicts the hierarchical clustering of interneurons within bat Pu. **H.** Bar plots showing the ratio of the number of *TAC3*, *PDGFD*, *PDGFD PTHLH PVALB-*, and *PDGFD PTHLH PVALB+* interneurons to the number of all interneurons across primate and non-primate CN, Pu, and C-Pu. **I.** Bar plots showing the ratio of the number of *CHAT*, *CCK VIP-*, *CCK VIP+* interneurons to the number of all interneurons across mouse caudoputamen, ferret CN, and bat, human, chimpanzee, rhesus macaque, and marmoset CN and Pu. Two tailed t-test was used to assess statistical significance between the groups. For comparisons where statistical significance was observed, p-values are highlighted in red. * indicates that using scCODA, the comparison is statistically significant and aligns with the direction of the results from the t-test. CN: Caudate Nucleus, Pu: Putamen, C-Pu: Caudoputamen. Error bars indicate mean ± standard error of the mean.

Surprisingly, we identified two interneuron types in bat that were either not found or found in very low abundance in the other species (**Fig. 4C and 4E**). These interneurons are characterized by the expression of transcription factors *LMO3* and co-expression of *FOXP2* and *TSHZ2* (**Fig. 4B**). Notably, both cell types are only present in bat Pu and not found in bat CN (**Fig. 4D**). Since *FOXP2* is also expressed in SPNs (**Fig. S3A**), we were curious about the specificity of the *FOXP2 TSHZ2 (BAT)* interneurons. We examined the expression of known SPN marker genes across these cells and found that these interneurons do not express SPN markers (**Fig. S4B**). We concluded that these cells are therefore interneurons found primarily in bats among the species we have examined.

To quantify this observation, we performed differential cellular abundance tests by using two-tailed t-tests (see **Methods**). We found that both *LMO3 (BAT)* and *FOXP2 TSHZ2 (BAT)* interneurons were significantly enriched in bat Pu samples compared to other species (**Fig. 4E** and **Fig S4C**). To test the validity of the t-test statistics, we again used scCODA,(Buttner, Ostner et al. 2021) with the following cell types as reference due to their high abundance and low dispersion (Buttner, Ostner et al. 2021): *PDGFD* interneurons for the CN and C-Pu tissues (**Fig. S4D**) and *SST NPY* interneurons for the Pu and C-Pu tissues (**Fig. S4E**). Relative to the proportional changes of the reference *SST NPY* interneurons, both the *LMO3 (BAT)* and the *FOXP2 TSHZ2 (BAT)* interneurons were significantly enriched in almost all comparisons between bat Pu and Pu of other species, except the comparison between bat Pu and marmoset Pu for *FOXP2 TSHZ2 (BAT)* interneurons (**Table S3** and **Fig. 4E**). Nonetheless, the scCODA results support the t-test findings (**Table S3** and **Fig. 4E)**.

The hierarchical clustering performed with all tissues and species showed *LMO3* and *FOXP2 TSHZ2* interneurons were close to *TH* and *TAC3* interneurons (**Fig. 4B**). These results indicate that in bat, *LMO3* and *FOXP2 TSHZ2* interneurons are cell types which may have been derived from a common thyrotropin expressing progenitor that also generates *TAC3* and *TH* interneurons. *LMO3* interneurons are 30% of all interneurons in bat Pu whereas *FOXP2 TSHZ2* interneurons comprise 9% of all interneurons in bat Pu (**Table S4**). Differential gene expression analysis showed that in *LMO3* interneurons, in addition to *LMO3,* the significantly upregulated genes include *ELAVL4*, *DCC*, and *KIF26B* whereas significantly downregulated genes include *RBFOX1*, *ERBB4*, and *NXPH1* (**Fig. 4F** and **Table S5**). Differential gene expression analysis in *FOXP2 TSHZ2* interneurons showed that in addition to *FOXP2*, *MEGF11, MEIS2*, and *MDIFC* genes were significantly upregulated whereas *GRIK2*, *RORA*, and *ERBB4* genes were significantly downregulated compared to other interneurons within bat Pu (**Fig. 4F**, **Table S6**). To characterize these interneurons better within bat Pu, we performed correlation analysis by generating a correlation matrix from the means of normalized gene expression values for each interneuron cell type (see **Methods**). The correlation analysis showed that *LMO3* interneurons were most similar to *FOXP2 EYA2* and *FOXP2 TSHZ2* interneurons (**Fig. 4G**). The *FOXP2 EYA2*-*FOXP2 TSHZ2 (BAT)-LMO3 (BAT)* clade was most similar to the *TH*-*TAC3* interneuron clade, indicating that these cells may have similar functions in bat Pu and validating the previous findings with all the species and tissues (**Fig. 4B** and **G**). These results suggest that by expanding single cell RNA sequencing analysis to different non-primate species we can identify new interneuron cell types.

### Identification of primate and human-specific interneurons

Recent work has found that *TH* and *TAC3* interneurons are transcriptomically similar to each other and concluded that *TH* interneurons are primarily present in mouse whereas *TAC3* interneurons are primarily present in primates (Krienen, Goldman et al. 2020, Garma, Harder et al. 2023, Corrigan, DeBerardine et al. 2024). In our dataset, in addition to reproducing the primate-specificity of *TAC3* interneurons compared to mouse (32% in CN and 23% in Pu, **Table S4,** p < 2.22 × 10^-6^ in CN and p = 9.5 10^-11^ in Pu, **Fig. 4H**), we also identified *TAC3* interneurons in bat (11% in CN, 5% in Pu **Fig. 4C** and **Table S4**). Thus, the inclusion of bat, a mammal with a distinct CN and Pu, demonstrates that this cell type is not primate specific. In our dataset, we also verified that *TH* interneurons are primarily present in mouse samples (16% of all interneurons, **Fig. S4C** and **Table S4**). However, we found that in addition to *TAC3* interneurons, bat samples contain *TH* interneurons as well (2% in Cu and 3% in Pu, **Fig. 4C** and **Table S4**), suggesting that *TH* interneurons are not specific to mouse. In bat Pu, we also found that *TH* and *TAC3* interneurons are correlated in terms of gene expression profiles, corroborating previous findings in other species (**Fig. 4G**) (Krienen, Goldman et al. 2020, Garma, Harder et al. 2023, Corrigan, DeBerardine et al. 2024). In our dataset, ferret samples did not contain *TH* interneurons, however, *TAC3* interneurons were present in ferret CN (7%, **Fig. 4C** and **Table S4**) as previously reported (Corrigan, DeBerardine et al. 2024). Together, these findings suggest that the *TAC3* and *TH* interneurons are conserved in adult striatum across approximately 94 million years of mammal evolution as previous studies have also shown (Schmitz, Sandoval et al. 2022, Corrigan, DeBerardine et al. 2024).

In addition to *TAC3* interneurons, we identified other interneuron subtypes containing a large number of cells: *PDGFD* interneuron classes (**Fig. 4A** and **Fig. 4SA**). In mouse it was previously found that *Pvalb* expression does not indicate a discrete class of striatal interneurons but highlights a striatal interneuron class expressing *Pthlh* (Munoz-Manchado, Bengtsson Gonzales et al. 2018). In *PVALB* interneurons, in addition to high *PVALB* expression, we also found high *PDGFD* expression, therefore we labeled these three classes of interneurons as: *PDGFD*, *PDGFD PTHLH PVALB-,* and *PDGFD PTHLH PVALB+* (**Fig. 4A-B** and **Fig. S4A**). We found that among these interneuron types, *PDGFD* and *PDGFD PTHLH PVALB-* were primarily present in primates (12% of all interneurons in CN, p = 2.3 × 10^-13^, and 12% of all interneurons in Pu, p = 9.3 × 10^-16^,and 16% of all interneurons in CN, p = 2.5 × 10^-7^, and 15% of all interneurons in Pu, p = 6.5 × 10^-10^, **Fig. 4C**, **Fig. 4H**, and **Table S4**) whereas *PDGFD PTHLH PVALB+* was present in CN and C-Pu of non-primates (in mouse C-Pu 31% of all interneurons, in ferret CN 28% of all interneurons, in bat CN 18% all interneurons, **Fig. 4C**, **Fig. 4H**, and **Table S4**). Interestingly, *PVALB* expressing *PDGFD PTHLH* interneurons are more abundant in non-primates (bat and ferret CN and mouse C-Pu) whereas the non-*PVALB* expressing counterpart of the same cell type is more abundant in primates (**Fig. 4H**). These findings highlight the importance of understanding the striatal *PVALB* activity in future comparative studies (primate vs non-primate) focused on physiology and function.

As our study is powered to interrogate human-specific changes by inclusion of novel chimpanzee CN and Pu datasets, we then investigated if there are any interneuron subtypes specifically altered in the human lineage in terms of proportional changes. We found *CHAT* interneurons are low in proportion in the human striatum, especially in CN (**Fig. 4I**). For both CN and Pu, *CCK VIP+* interneurons are significantly higher in proportion in humans compared to other species although there is a trend but no significance when human and chimpanzee *CCK VIP+* interneuron proportions were compared (**Fig. 4I**). In CN, performing scCODA and using *SST NPY* interneurons as a reference cell type, we could only validate the t-test statistics results of the differential cellular abundance of the *CHAT* interneurons for comparisons of human-mouse and human-bat (**Table S7** and **Fig. 4I**). These results suggest that within primates, *CHAT* interneuron proportion is similar and this is reduced compared to non-primate CN and mouse C-Pu. Similarly, using the same tissue (CN) and the reference cell type (*SST NPY*), scCODA results for *CCK VIP+* interneurons validated the t-test results only in the human-marmoset comparison (**Table S7 and Fig. 4I**). In the human-mouse comparison, although the log_2_fold change (−0.86) was lower than in the human-marmoset comparison (log_2_FC = −0.8), the algorithm considered it not significant (−0.734 - 0.029, HDI 3% - HDI 97%) (**Table S7 and Fig. 4I**). In Pu, compared to the reference cell type *PDGFD*, scCODA yielded significantly higher abundance for *CCK VIP+* interneurons in human only compared to mouse C-Pu (**Table S8 and Fig. 4I**). Our findings highlight species-specific differences in the composition of striatal interneuron cell types, with human samples showing distinct patterns compared to mouse and marmoset samples. However, the Bayesian approach may require a larger sample size to achieve statistical significance when comparing with other species.

## Discussion

Cell type-specific genomics of subcortical brain regions remain underexplored, with a limited availability of single-cell resolution high-throughput datasets for these areas. Dorsal striatum is an integral part of the forebrain and is involved in decision making, motor control, and reward processing (Balleine, Delgado et al. 2007). In this study, we generated and analyzed a comprehensive snRNA-seq dataset, encompassing the CN and Pu across four primates—human, chimpanzee, rhesus macaque, and common marmoset—and three non-primate species: pale spear-nosed bat, ferret, and mouse C-Pu (**Fig. 1**). By including bat CN and Pu, we were able to identify novel interneuron cell types, *LMO3* and *FOXP2 TSHZ2* in the bat Pu, whose functional roles will be explored in future studies. These findings help underscore the unique characteristics of non-primate striatal anatomy and physiology (**Fig. 4A-E**). Additionally, we identified significant differences in primates, including lower neuron-to-glia ratios reflecting the allometric scaling of the neuron density and relative glial cell density invariance across different brain sizes as previously shown in ventral striatum (Hirter, Miller et al. 2021) and other regions (Herculano-Houzel 2014) (**Fig. 2**) and higher eSPN-to-SPN ratios in comparison to non-primate striata (**Fig. 3**). Furthermore, we could also identify tissue specific differences. In bat CN, we found there is a higher neuron-to-glia ratio than in bat Pu. The ratios of *PDGFD PTHLH PVALB+* interneurons to all interneurons were higher in non-primate CN samples compared to primate CN samples; however, bat Pu showed similar proportions to Pu of primates (**Table S4**). Our work parallels previous work on striatum by highlighting the conservation of important cell types and transcriptomic profiles across approximately 94 million years of mammal evolution (Gokce, Stanley et al. 2016, Munoz-Manchado, Bengtsson Gonzales et al. 2018, Saunders, Macosko et al. 2018, Krienen, Goldman et al. 2020, He, Kleyman et al. 2021). Together, our results suggest that the species-specific and tissue-specific specializations of interneuron cell types and cell type abundances could offer insights into species-specific behavioral differences.

Although they are few in number, interneurons not only regulate the basal ganglia circuitry by integrating incoming signals from various brain regions and influencing the activity of SPNs to modulate output signals, but they are also involved in changes related to plasticity and adaptation in disease conditions (Tepper, Tecuapetla et al. 2010, Conti, Chambers et al. 2018, Mallet, Leblois et al. 2019). Previous studies reported differences in striatal interneuron number, location and physiological features across various species (Graveland and DiFiglia 1985, Bernacer, Prensa et al. 2012, Petryszyn, Beaulieu et al. 2014, Lecumberri, Lopez-Janeiro et al. 2018, Munoz-Manchado, Bengtsson Gonzales et al. 2018, Krienen, Goldman et al. 2020). The presence of the interneuron cell types, predominantly observed in pale spear-nosed bat, underscores fundamental changes in the species-specific evolution of the striatum. Hierarchical clustering analysis indicates that the newly identified *LMO3* and *FOXP2 TSHZ2* interneurons share significant transcriptomic similarities with the *TAC3*-*TH* interneurons, indicating that they may originate from the same class of progenitors as the *TAC3*-*TH* interneurons (**Fig. 4B**). Previous studies have shown the extensive heterogeneity in tyrosine hydroxylase (TH) expressing cells of adult mice (Ibanez-Sandoval, Tecuapetla et al. 2010, Unal, Ibanez-Sandoval et al. 2011). In embryonic mouse stem cells, *Lmo3*, together with *Pou3f4*, was found to be necessary for the differentiation of cortical inhibitory neurons (Au, Ahmed et al. 2013). Additionally, during both human and mouse dopaminergic neuron development, it was found that *LMO3* was co-expressed by a subset of TH cells (La Manno, Gyllborg et al. 2016). However, further studies are needed to understand the function of *LMO3* in bat adult striatal interneurons. Previous studies suggest that FOXP2 has a role in striatal function and speech related motor skills (Vargha-Khadem, Watkins et al. 1998, Watkins, Vargha-Khadem et al. 2002, Liegeois, Baldeweg et al. 2003, French, Jin et al. 2012). In adult mice, *Tshz2* was shown as a marker for a cell type found in prefrontal cortex (Lui, Nguyen et al. 2021) as well as amygdala excitatory neurons (Sun, Liu et al. 2024); however, the function of *TSHZ2* has yet to be characterized in adult striatal interneurons and in other species. Additionally, it is not yet understood whether these novel interneurons emerge from the same progenitor cell as *TH*-*TAC3* interneurons. Future electrophysiological and imaging studies can delineate the functions of these novel interneuron cell types in bat Pu.

With the advancement of single-cell technologies, cell type innovation has been discovered across species in brain (Saunders, Macosko et al. 2018, Krienen, Goldman et al. 2020, Caglayan, Ayhan et al. 2023) and in other organs (Mah and Dunn 2024, Williamson, Liyanage et al. 2024). Expanding evolutionary distance by incorporating a diverse range of species and tissues should uncover alterations in cell types and proportions and enhance our understanding of species-specific variation in cellular identity and behavior across evolutionary lineages. As an example of including non-model species to investigate cell type function, in a recent study performed on oldfield mice (*Peromyscus polionotus*), a link between a recently evolved-cell type and parental behavior was found (Niepoth, Merritt et al. 2024). In summary, our findings corroborate previous snRNA-seq studies on striatum and highlight the critical importance of incorporating a broad range of species when identifying novel cell types and investigating variations in cellular abundance. Future studies examining the functional roles of these newly identified cell types will enhance our understanding of subcortical brain regions and may provide insights into species-specific behavior.

## Materials & Methods

### Tissue collection

#### Human and non-human primates

All human tissue was obtained from NIH NeuroBioBank. Chimpanzee tissues were obtained from individuals previously housed at either the Emory National Primate Research Center or the Michale E. Keeling Center for Comparative Medicine and Research. Macaque tissues were obtained from the Michale E. Keeling Center for Comparative Medicine and Research. Caudate and putamen were dissected from frozen postmortem tissue slabs (**Table S1**).

#### Bats

The adult *Phyllostomus discolor* bats originated from a breeding colony situated in the Department Biology II of the Ludwig-Maximilian University of Munich or in St Mary’s Animal Unit of the University of St Andrews. Animals were kept under semi-natural conditions (12 h day/12 h night cycle, 65%–70% relative humidity, 28°C) with access to food and water ad libitum. All the experiments complied with the principles of laboratory animal care and the regulations of the current version of the German Law on Animal Protection and the Animal Scientific Procedures Act (1986) under Home Office (UK) supervision. The number of animals used in terminal experiments was reported to the Munich veterinary office and the Home Office (UK).

In order to collect the samples used in the single nuclear RNA sequencing experiments, the animals were euthanized by an intraperitoneally applied lethal dose of pentobarbital (0.16 mg/g bodyweight) followed by decapitation. The brain was extracted, embedded in 3% low melting point agarose and sliced in 500 µm coronal slices on a vibratome (Precisionary Compresstome® VF-200 and Leica VT1200S) in ice-cold buffer solution (3 mM KCl, 3 mM MgCl2.6H2O, 23 mM NaHCO3, 1.2 mM NaH2PO4.H2O, 210 mM Sucrose, 11 mM Glucose). Tissue punches (1.5 mm diameter, KAI Europe GmbH) were taken from the slices and were snap-frozen in liquid nitrogen.

To harvest tissue used in Immunofluorescence experiments, the animals were sedated with MMF (medetomidine, midazolam and fentanyl at a dosage of 0.4, 4.0 and 0.04 µg/g body weight, respectively), euthanised with an intraperitoneal injection of pentobarbital (0.16 mg/g bodyweight) and perfused transcardially with approximately 200 ml of phosphate-buffered saline (PBS; pH 7.4; 0.9% NaCl and 1% heparin 25.000 IU) for 10 minutes, followed by approximately 200 ml of 4% paraformaldehyde in PBS over ∼15-20 minutes. Samples were embedded in OCT (Optimal Cutting Temperature) compound and frozen by isopentane bath. The samples are left in dry ice and stored at −80⁰C until sliced. The brains were sliced from a coronal position, using a Leica CM1860 cryostat, 15 microns thick and placed on positively charged glass slides (Superfrost plus Gold). The slides were then stored at −80⁰C until used.

### Single-nucleus RNA library generation and sequencing

#### Bat samples

Nuclei for snRNA-seq were isolated from bat putamen and caudate brain tissue. Briefly, the tissue was homogenized in 500 µL of ice-cold Nuclei EZ lysis buffer (EZ PREP NUC-101, Sigma) using a pre-chilled glass Dounce homogenizer. Debris was removed via density gradient centrifugation using Nuclei PURE 2 M sucrose cushion solution and Nuclei PURE sucrose cushion buffer (Nuclei PURE prep isolation kit, no. NUC201-1KT, Sigma Aldrich). These two solutions were mixed in a 9:1 ratio, and 500 µL of the sucrose solution was added to a 2-ml Eppendorf tube. The homogenized nuclei suspension was mixed with 900 µL of sucrose cushion solution by pipetting 10 times, resulting in a 1,400 µL volume, which was carefully layered over the sucrose buffer without mixing. The samples were centrifuged at 13,000 x g for 45 min at 4 °C, and all but 100 µL of supernatant was discarded. Nuclei were resuspended in 300 µL of nucleus suspension buffer (NSB), consisting of 1× PBS, 2% BSA (no. AM2618, Thermo Fisher Scientific), and 0.4 U µL −1 RNAse inhibitor (no. AM2694, Thermo Fisher Scientific). Samples were then centrifuged at 550 x g for 5 min at 4 °C, and all but 50 µL of supernatant was discarded. The nuclear pellet was resuspended in NSB and filtered through a 40-μm Flowmi cell strainer (Bel-Art Products, #H13680-0040) into a new tube. The nuclei concentration was measured using Hoechst and decreased to 1,000 nuclei per µL with NSB if necessary.

Droplet-based snRNA-seq libraries were prepared using the Chromium Single Cell 3′ v3.1 (10x Genomics) kit, following the manufacturer’s protocol. Libraries were sequenced on an Illumina NovaSeq 6000.

#### Human, chimpanzee, and rhesus macaque samples

Nuclei for snRNA-seq were isolated from the putamen and caudate nuclei of human, chimpanzee, and rhesus macaque tissues. Briefly, the tissue was homogenized in 750 µL of ice-cold NP40 lysis buffer (10 mM Tris-HCl pH 7.4, 10 mM NaCl, 3 mM MgCl₂, 0.1% Nonidet P40 Substitute, 1 mM DTT, 0.2 U/µL RNAse inhibitor) using a pre-chilled glass Dounce homogenizer. An additional 750 µL of ice-cold NP40 lysis buffer was added, and the mixture was pipette-mixed during a 5-minute incubation on ice. Nuclei were centrifuged at 500 x g for 5 minutes at 4°C, resuspended in 1 mL of ice-cold lysis buffer, and incubated on ice for 5 minutes with gentle pipette mixing throughout. The nuclei were then centrifuged again at 500 x g for 5 minutes at 4°C, and all but 50 µL of supernatant was discarded. Next, 500 µL of nucleus suspension buffer (NSB) containing 1× PBS, 1% Ultrapure BSA (no. AM2618, Thermo Fisher Scientific), and 0.2 U/µL RNAse inhibitor (no. AM2694, Thermo Fisher Scientific) was added, and the sample was incubated for 5 minutes on ice without mixing. The pellet was resuspended, centrifuged again at 500 x g for 5 minutes at 4°C, and all but 50 µL of supernatant was discarded. The nuclei were then resuspended in 500 µL of NSB and filtered through a 70-μm Flowmi cell strainer (no. H13680-0070, Bel-Art) into a 1.5 mL tube.

Debris was removed using density gradient centrifugation with Nuclei PURE 2 M sucrose cushion solution and Nuclei PURE sucrose cushion buffer (Nuclei PURE prep isolation kit, no. NUC201-1KT, Sigma Aldrich). The solutions were mixed in a 9:1 ratio, and 500 µL of the resulting sucrose solution was added to a 2-ml Eppendorf tube. A 900 µL volume of sucrose buffer was added to 500 µL of isolated nuclei in NSB, and the resulting 1,400 µL suspension was layered on top of the sucrose buffer. The gradient was centrifuged at 13,000 x g for 45 minutes at 4°C. After centrifugation, all but 100 µL of supernatant was removed, and the nuclei were resuspended in 400 µL of NSB, centrifuged at 500 x g for 5 minutes at 4°C, and all but 50 µL of supernatant was discarded. The nuclei were then resuspended in 200 µL of NSB and filtered through a 40-μm Flowmi cell strainer (no. H13680-0040, Bel-Art). The concentration of nuclei was determined using 0.4% trypan blue (no. 15250061, Thermo Fisher Scientific) and adjusted to a final concentration of 1,000 nuclei per µL with NSB.

Droplet-based snRNA-seq libraries were prepared using Chromium Single Cell 3′ v3.1 (1000121, 10x Genomics) according to the manufacturer’s protocol. Libraries were sequenced on an Illumina NovaSeq 6000.

### Single nucleus RNA sequencing dataset alignment and quality control

We obtained raw snRNA-seq data for human, chimpanzee, macaque, and pale spear-nosed bat libraries from the McDermott Sequencing Core at UT Southwestern in binary base call (BCL) files. The BCL files were demultiplexed using CellRanger bcl2fastq v2.20.0 and CellRanger mkfastq (10x Genomics Cell Ranger 6.0.0) with default parameters to generate FASTQ files. For the previously published rhesus macaque (He, Kleyman et al. 2021), common marmoset (Lin, Kelly et al. 2022, Krienen, Levandowski et al. 2023), mouse (Krienen, Goldman et al. 2020), and ferret (Krienen, Goldman et al. 2020) datasets, the FASTQ files were downloaded and preprocessing was carried out similarly to our own datasets.

Since any non-human primate has a less accurate GTF file than human GTF file, we converted the positions of every non-human primate to the human annotation file using liftoff (v1) (Shumate and Salzberg 2021). We used the following genomes: Chimpanzee (*Pan troglodytes*) panTro5, macaque (*Macaca mulatta*) rheMac10, common marmoset (*Callithrix jacchus*) caljac3, mouse (*Mus musculus*) mm10, pale spear-nosed bat (*Phyllostomus discolor*) GCF_004126475.2_mPhyDis1.pri.v3, and ferret (*Mustela putorius furo*) GCF_011764305.1_ASM1176430v1.1. The index for each genome file was generated with CellRanger mkref (10x Genomics Cell Ranger 6.0.0). The FASTQ files were aligned to their corresponding genomes using CellRanger count (10x Genomics Cell Ranger 6.0.0).

To remove ambient RNA contamination, CellBender (Fleming, Chaffin et al. 2023) was used on the raw count matrices. With these filtered gene-cell matrices the Seurat objects were generated. To assess the health of the nuclei, we used intronic read ratios (Caglayan, Liu et al. 2022). Briefly, we counted the number of reads that are mapped to introns and divided them to the total number of mapped reads within a cell. During clustering, we removed the clusters with exceptionally low mean intronic read ratios (< 0.5). In our dataset, the healthy nuclei had intronic read ratios ∼ ≥ 0.7 (**Fig. S1B**).

### Single nucleus RNA sequencing cell type annotation and cellular compositional abundance analysis

In addition to clusters with low intronic read ratios, we also removed clusters with an unusually high number of detected genes accompanied with a high level of expression of at least two typically distinct marker genes as potential nuclear doublets. In addition to doublets, we also removed non-cells (empty), endothelial cells, and pericytes (*FLT1*, *DUSP1*, *EBF1, PDGFRB* expressing clusters) from the dataset. We re-clustered the nuclei and repeated this process if needed until no such clusters were found. We then used canonical marker genes (for example, *GAD1* for inhibitory neurons) (Munoz-Manchado, Bengtsson Gonzales et al. 2018, Martin, Calvigioni et al. 2019, Khrameeva, Kurochkin et al. 2020, Krienen, Goldman et al. 2020) to broadly annotate striatal nuclei in each species. Major cell types were defined as: SPNs, interneurons, astrocytes, mature oligodendrocytes (MOLs), oligodendrocyte progenitor cells (OPCs), and microglia.

After broad annotation, we extracted each broad category (for example, SPNs) from all species and integrated them using the default approach in Seurat v3 across all samples (SelectIntegrationFeatures, PrepSCTIntegration, FindIntegrationAnchors). During SPN filtering and annotation, we removed samples with less than 300 dSPN or iSPNs (human sample: Sample 242939, macaque samples SRR13808459, SRR13808461, SRR13808466, and SRR1380847 were removed). We annotated the SPN types and matrix and striosome clusters based on the canonical markers (Gokce, Stanley et al. 2016, Martin, Calvigioni et al. 2019). During interneuron filtering and annotation, we removed clusters which were not consistently present across all the groups within each species-tissue pair. We annotated the interneuron cell types based on the canonical markers (Munoz-Manchado, Bengtsson Gonzales et al. 2018, Krienen, Goldman et al. 2020). For both SPN and interneuron integrations, only protein coding genes which were orthologous across all the species were used. The orthologous genes were obtained using NCBI Datasets tool (v 16.35.1).

Cellular proportions were calculated as dividing the number of neurons (SPNs and interneurons) to the total number of cells (SPNs, interneurons, and glia) within each sample. The mean proportions, standard deviations, sample numbers, and standard errors of mean for each tissue-species pair are provided (**Table S2**). These proportions were compared using a two-tailed t-test, with p-values ≤ 0.05 considered statistically significant.

Cellular proportions of each cell type within SPNs or interneurons were calculated (**Table S4**). These proportions were compared using a two-tailed t-test, with p-values ≤ 0.05 considered statistically significant. To better accommodate the inherent characteristics of proportional data and account for different underlying assumptions, an alternative approach was employed utilizing a Bayesian model with scCODA (Buttner, Ostner et al. 2021), where a false discovery rate (FDR) threshold of ≤ 0.1 was applied.

### Differentially expressed gene (DEG) and correlation analysis

When finding the conserved eSPN markers, we performed DEG analysis using FindMarkers from the Seurat package (Wilcoxon rank sum test) by comparing eSPN and other cells. To remove the neuron specific DEGs and only focus on the eSPN-specific ones, we additionally performed DEG analysis by running FindMarkers using eSPN and other neurons (dSPNs, iSPNs, and interneurons). We then found the intersection of these lists of DEGs to find the eSPN-specific DEGs for that species. We repeated this process for all species. We then got the intersection of the eSPN-DEGs for all the species and filtered these DEGs by keeping the genes with scaled average expression greater than 1 for eSPN and plotted in dot plot (**Fig. 3F**).

Hierarchical clustering was performed with the BuildClusterTree function of Seurat v3 with the default parameters: The centroids (mean values of the principal components) for each cluster were calculated to reduce the dataset to one representative vector per cluster. The pairwise Euclidean distances between the clusters were calculated and using Ward D2 method to minimize the variance within each cluster, the distance matrix was created. This distance matrix was visualized using PlotClusterTree function.

Similarly to eSPN DEGs, we used FindMarkers from Seurat (Wilcoxon rank sum test) to determine the *LMO3* and *FOXP2 TSHZ2* interneuron specific DEGs within bat putamen compared to all other interneurons and showed the statistically upregulated (log2 fold change > 0.6 adjusted p value < 0.05) and downregulated (log2 fold change < −0.6 adjusted p value < 0.05) genes in the volcano plots (**Fig. 4F**).

Pearson correlation analysis with default parameters was performed to assess the similarity between gene expression profiles of D1-D2 hybrid cells from the macaque dataset (He, Kleyman et al. 2021) and our SPN dataset as well as interneuron cell types within bat Pu. The resulting correlation values were visualized with a heatmap (**Fig. S3G** and **Fig. 4G**).

### Single molecule fluorescence in situ hybridization (smFISH)

smFISH was performed using RNAScope Multiplex v2 Fluorescent assay. All steps were performed according to the manufacturer’s instructions (fresh frozen tissues for human and fixed frozen tissues for bats and mice) but with the addition of Sudan Black B. Sudan Black B (0.05%) added to the tissues after application of DAPI (Advanced Cell Diagnostics (ACD) #320858) to quench autofluorescence. We used the following: Bat *MOG* probe (ACD #1076431-C2) with opal dye 570 (Akoya Biosciences #NC1601878), human *MOG* (ACD #543181-C2) with opal dye 570 (Akoya Biosciences #NC1601878), bat *RBFOX3* probe (ACD #1279941-C3) with opal dye 690 (Akoya Biosciences #NC1605064), human *RBFOX3* probe (ACD #415591-C3) with opal dye 690 (Akoya Biosciences #NC1605064), human *DRD1* (ACD #524991-C2) with opal dye 570 (Akoya Biosciences #NC1601878), human *DRD2* (ACD #553991-C3) with opal dye 520 (Akoya Biosciences #NC1601877), human *CASZ1* (ACD #882211-C1) with opal dye 690 (Akoya Biosciences #NC1605064), mouse *Drd1* (ACD #461901-C1) with opal dye 570 (Akoya Biosciences #NC1601878), mouse *Drd2* (ACD #406501-C2) with opal dye 520 (Akoya Biosciences #NC1601877), and mouse *Casz1* (ACD #502461-C3) with opal dye 690 (Akoya Biosciences #NC1605064).

Imaging *RBFOX3* and *MOG* was performed at ×20 magnification (zoom 2.5) and imaging *DRD1*, *DRD2*, and *CASZ1* was performed at ×20 magnification (zoom 1 and zoom 2.5) using a Zeiss LSM 880 in in the UT Southwestern Neuroscience Microscopy Facility.

Cell numbers (*MOG*+, *RBFOX3*+, etc.) were quantified manually in double blind manner using Fiji. For example, MOG was defined as double positive cells for *MOG* and DAPI, RBFOX3 was defined as double positive cells for *RBFOX3* and DAPI.

## Code and Data Availability

The human, chimpanzee, and pale spear-nosed bat dorsal striatum snRNA-seq datasets were deposited to GEO (accession number GSE293075). The following published dorsal striatum snRNA-seq datasets were downloaded from GEO and used in the analysis: Rhesus macaque dataset with accession number GSE167920, marmoset datasets with accession numbers GSE151761 and GSE165578, mouse and ferret datasets with accession number GSE151761.

All the scripts used in the study were deposited to https://github.com/konopkalab/Comparative_striatum

## Acknowledgements

We thank Amber Main and Siddhartha Lavu for technical support. We thank Prof. Lutz Wiegrebe^†^ and Dr. Uwe Firzlaff for their support with bat tissue collection. Chimpanzee tissues were provided by the National Chimpanzee Brain Resource which was supported by NINDS (R24NS092988). Human tissues were obtained from the NIH NeuroBioBank. Macaque tissues were obtained from Michale E. Keeling Center for Comparative Medicine and Research. G.K. is a Jon Heighten Scholar in Autism Research and Townsend Distinguished Chair in Research on Autism Spectrum Disorders at UT Southwestern. This work was partially supported by the James S. McDonnell Foundation 21st Century Science Initiative in Understanding Human Cognition Scholar Award (220020467), NHGRI (HG011641), NINDS (NS115821, NS126143) and NIMH (MH134809, MH126481, MH103517) to G.K, and NSF (EF-2021785, DRL-2219759), NHGRI (HG011641), and NIMH (MH134809) to C.C.S. This project has received funding from the European Research Council (ERC) under the European Union’s Horizon 2020 research and innovation programme (ERC Consolidator Grant, agreement: BATSPEAK and 101001702) and a UKRI Future Leaders Fellowship (MR/T021985/1), both awarded to SCV.

---

**Fig. S1.**
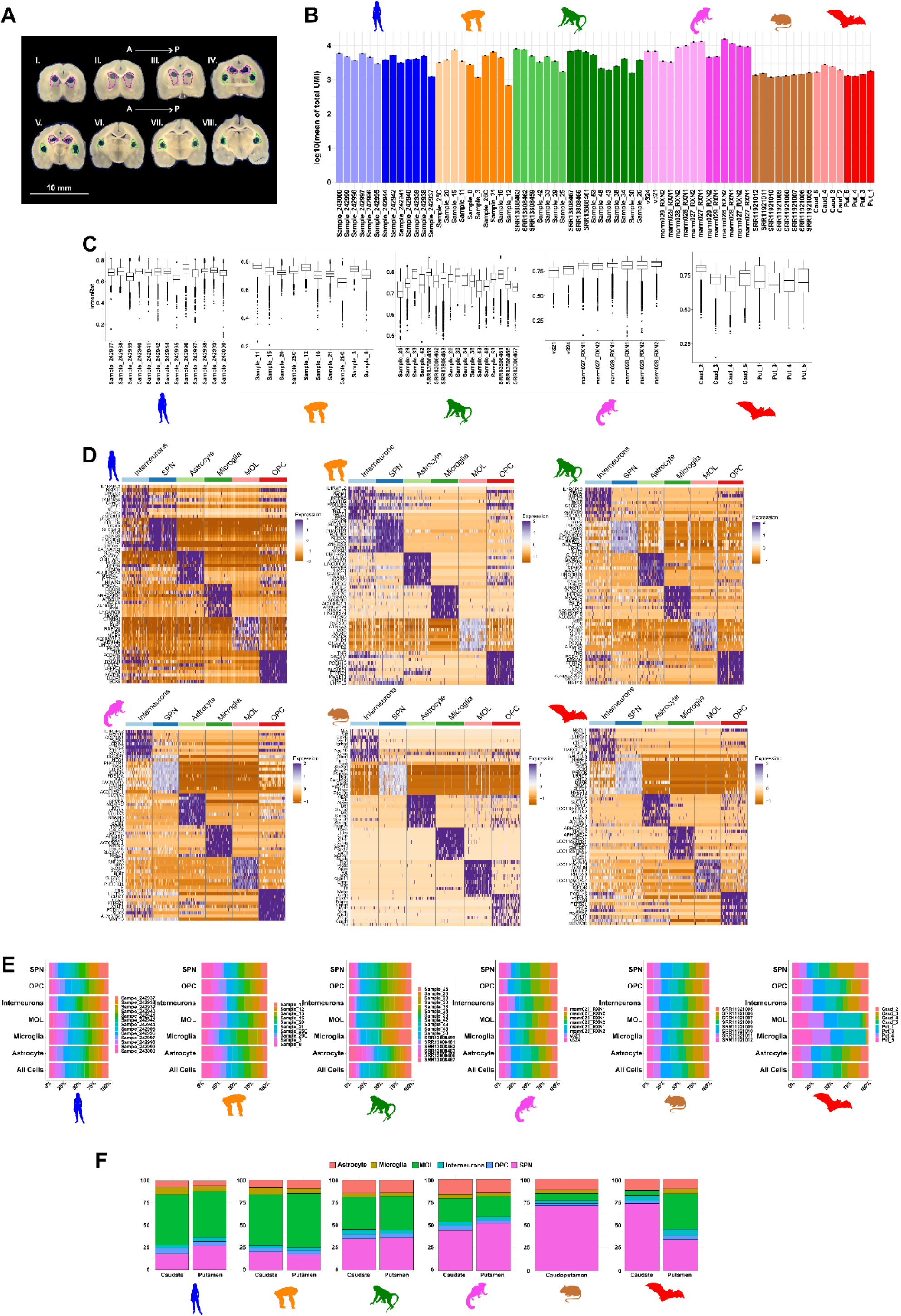
Quality control metrics for major cell type annotations across species. **A.** Representative photos of bat brain tissue dissection. Punches were made from the coronal sections of bat CN and Pu. Drawings in magenta indicate the punches made through the CN (I-IV) and drawings in green indicate the punches made through the Pu (IV-VIII). The scale bar marks 10 mm. **B.** Bar plots illustrate the mean total number of unique molecular identifiers (UMIs) per cell across samples, displayed on a logarithmic scale (base 10), segmented by species and tissue. The light colors represent the CN and the dark colors represent either Pu or C-Pu. **C.** Box plots depicting the intronic read ratios for samples, across all species. **D.** Heatmap depicting the expression levels of the top 10 most highly differentially expressed genes within each major cell type annotation in each species. **E.** Cell type composition of each sample across species. **F.** Bar plots displaying the proportional composition of major cell type annotations within each tissue type (CN, Pu, caudoputamen) and species. CN: caudate nucleus, Pu: putamen. A: anterior. P: posterior.

**Fig. S2.**
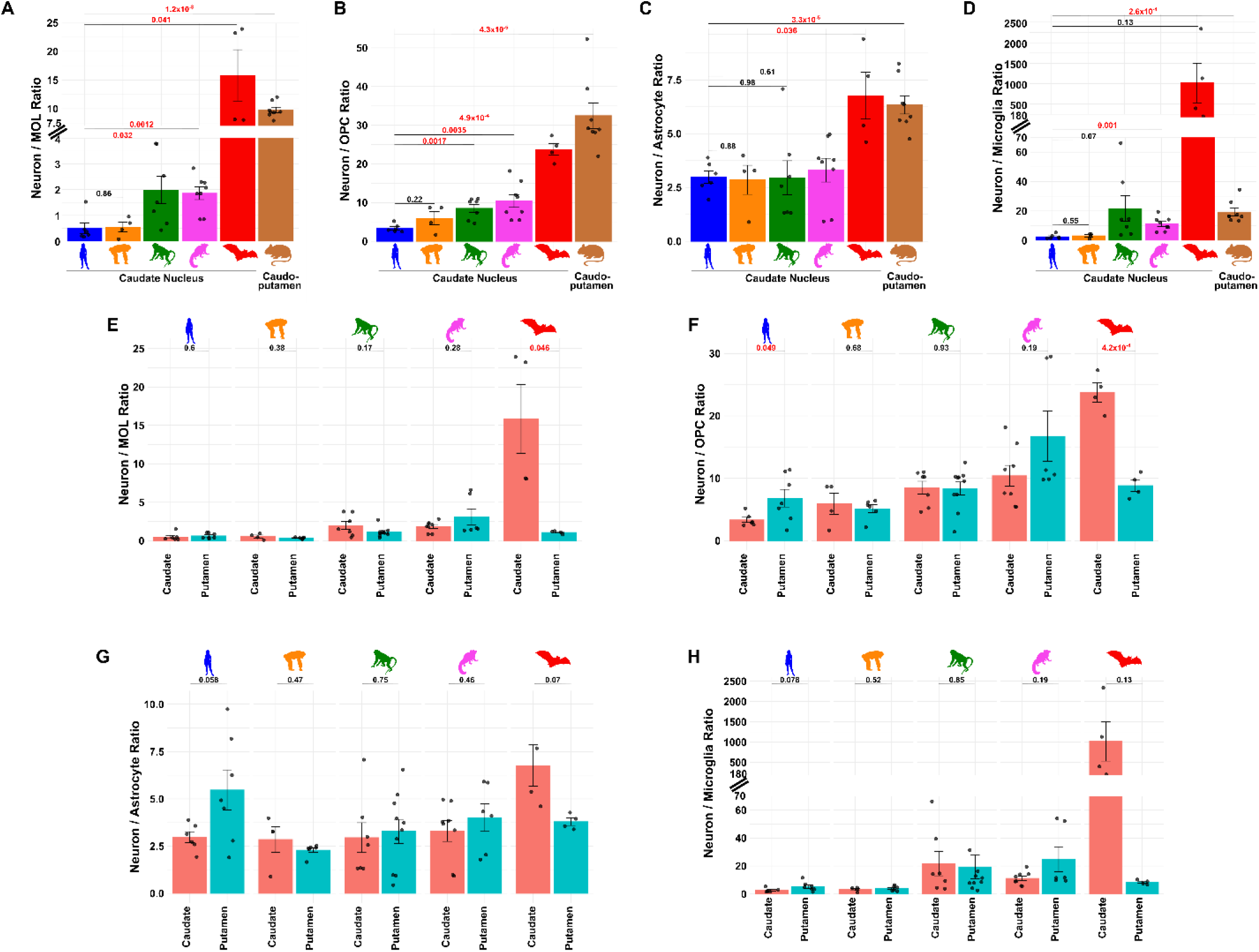
Ratios of neuron to subtypes of glia. **A-D.** Bar plots illustrating the ratios of neuron counts to counts of various glial subtypes: **A.** Mature oligodendrocytes (MOL), **B.** Oligodendrocyte progenitor cells (OPC), **C.** Astrocytes, and **D.** Microglia. These ratios are depicted for the caudate nucleus (CN) across all species, and the caudoputamen in mouse. **E-H.** Bar plots displaying the ratios of neuron counts to counts of glial subtypes in the following categories: **E.** Mature oligodendrocytes (MOL), **F.** Oligodendrocyte progenitor cells (OPC), **G.** Astrocytes, and **H.** Microglia. These ratios are shown for both the CN and putamen (Pu) across all species. Two tailed t-test was used to assess statistical significance between the groups. For comparisons where statistical significance was observed, p-values are highlighted in red. Error bars indicate mean ± standard error of the mean.

**Fig. S3.**
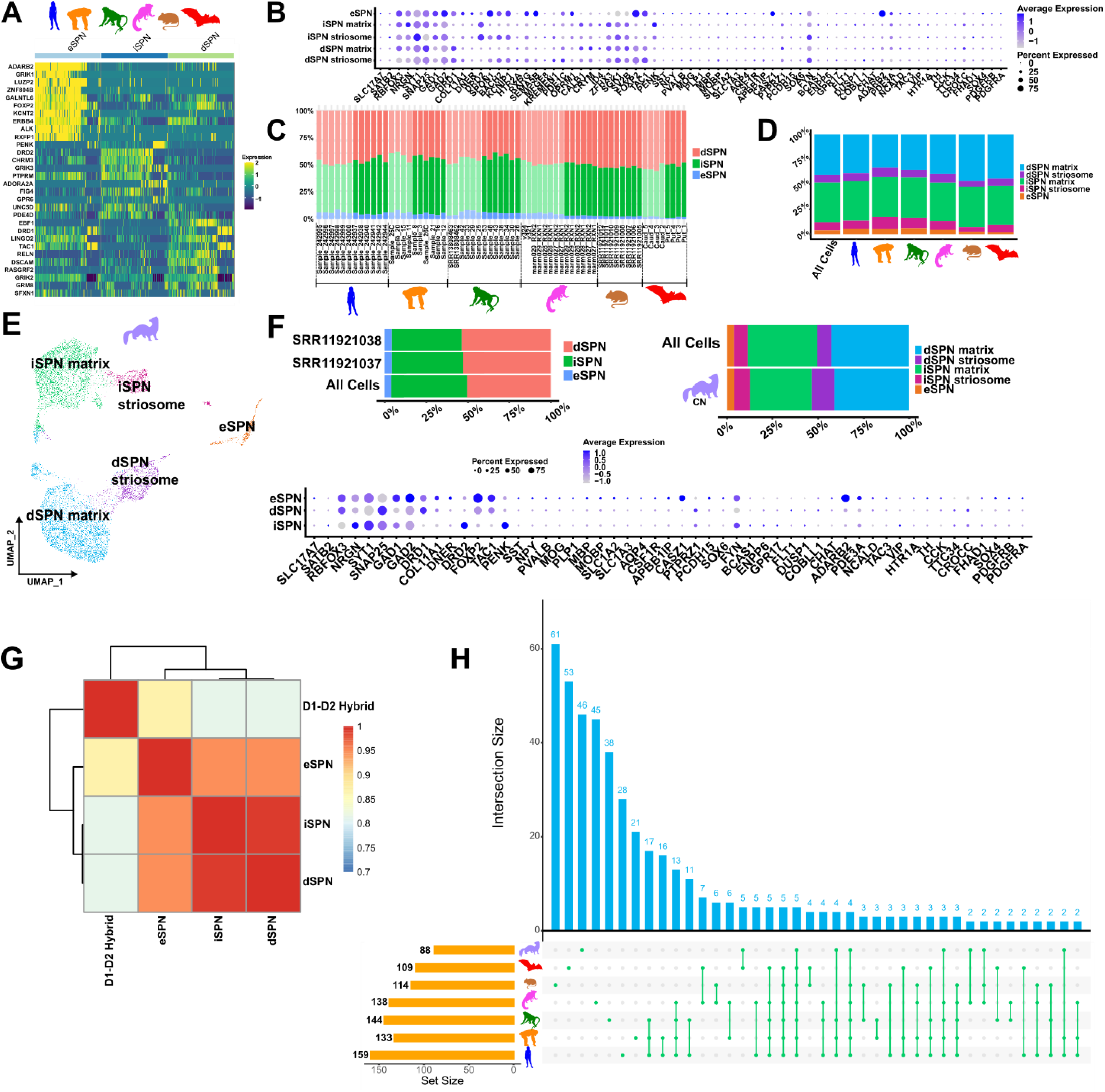
Spiny projection neuron (SPN) characteristics. **A.** Heatmap showing the transcriptomic profile of the top 10 genes in each SPN subtype: dSPN, iSPN, and eSPN across human, chimpanzee, rhesus macaque, marmoset, mouse, and bat. **B.** Dot plot of marker genes used to determine the SPN subtypes: dSPN striosome, dSPN matrix, iSPN striosome, iSPN matrix, and eSPN. **C.** Stacked bar plot depicting the percentages of each SPN subtype (dSPN, iSPN, and eSPN) across samples. The samples were grouped together according to the species they belong to. The light colors represent the CN and the dark colors represent either Pu or C-Pu. **D.** Cellular composition of SPN subtypes: dSPN striosome, dSPN matrix, iSPN striosome, iSPN matrix, and eSPN across human, chimpanzee, rhesus macaque, marmoset, mouse, and bat. **E.** UMAP of SPNs of ferret CN (n=2). **F.** The stacked bar plot of SPN subtypes: dSPN, iSPN, and eSPN across ferret samples and the cellular composition of SPN subtypes: dSPN striosome, dSPN matrix, iSPN striosome, iSPN matrix, and eSPN in ferret dataset. Dot plot of the marker genes in the ferret dataset. **G.** Heatmap showing the Pearson correlation matrix between the normalized gene counts of the D1-D2 hybrid cells in the published macaque dataset (He, Kleyman et al. 2021) and our SPN dataset. The dendrogram depicts the hierarchical clustering of these cells. **-H.** Upset plot of differentially expressed genes (DEGs) in eSPNs compared to other cell types across species: human, chimpanzee, rhesus macaque, marmoset, mouse, bat, and ferret. Each blue bar indicates the number of common DEGs for the indicated combination of species in green. Yellow bars indicate the total number of eSPN DEGs found in each species. Two tailed t-test was used to assess statistical significance between the groups. For comparisons where statistical significance was observed, p-values are highlighted in red. CN: Caudate nucleus and Pu: putamen.

**Fig. S4.**
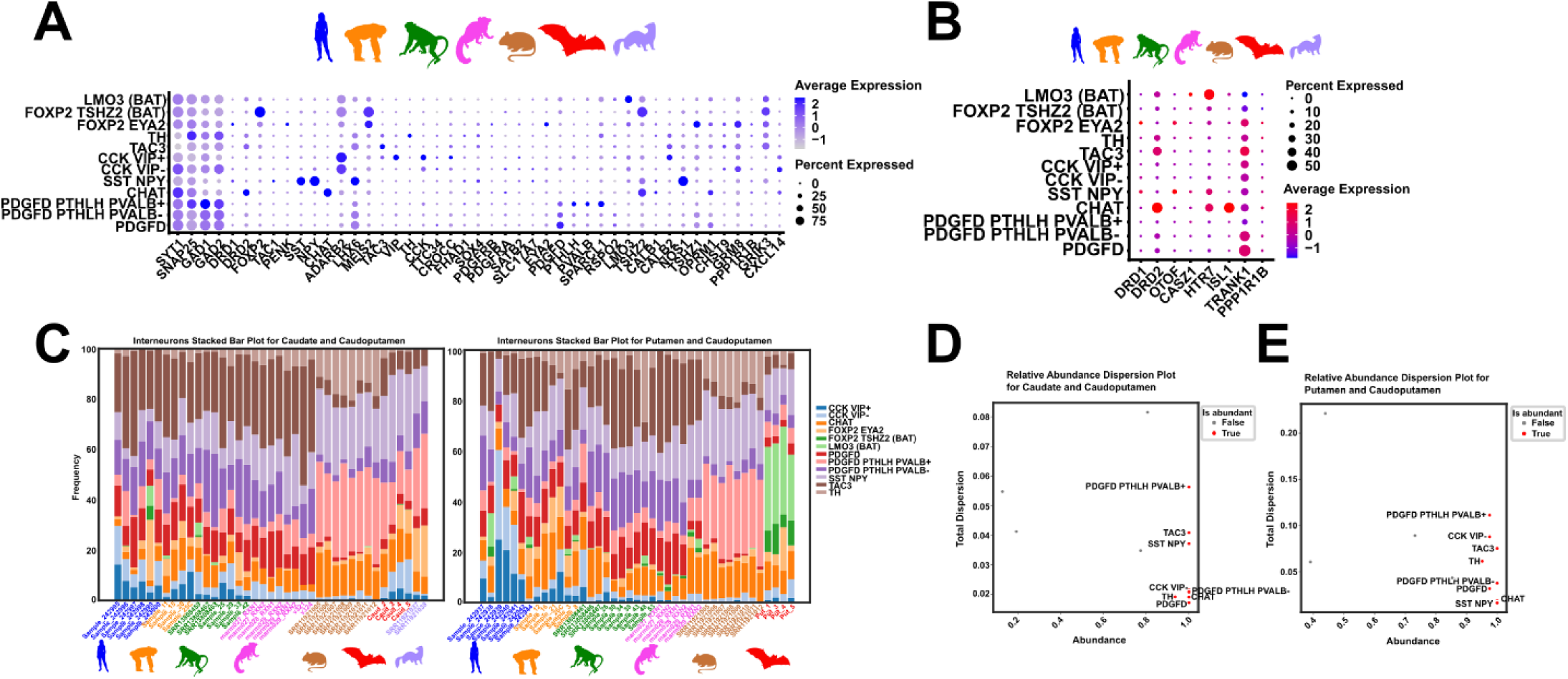
Striatal interneurons across all species. **A.** Dot plot showing the expression of marker genes across striatal interneuron cell types of human, chimpanzee, rhesus macaque, marmoset, mouse, bat, and ferret. **B.** Dot plot showing the expressions of SPN marker genes across striatal interneuron cell types of human, chimpanzee, rhesus macaque, marmoset, mouse, bat, and ferret. **C.** Sample level distribution of striatal interneuron cell types in CN and C-Pu and in Pu and C-Pu. **D-E.** Dispersion and cellular abundance plots for **D.** CN and **E.** Pu. The striatal interneuron cell types indicated in red were found to be higher than 0.9 in abundance. CN: Caudate Nucleus, Pu: Putamen, C-Pu: Caudoputamen.

